## Supplementary Materials for "Dynamics of history-dependent perceptual judgment"

Supplementary Figures:

**S1.** Gradual psychometric shift, excluding trials with incorrect *n-1* choice, suggests that *n-1* Δ*speed* drives the repulsive bias.

**S2.** The effect of stimulus *n-1* on psychometric parameters and classification of stimulus *n* indicates horizontal psychometric shift.

**S3.** Discrete history model improves prediction about judgment of ambiguous stimuli only.

**S4.** Stimulus-dependent bias builds up between trials also when removing incorrect trials and considering only early session trials.

**S5.** Vibrotactile categorization task transferred to human tactile perception.

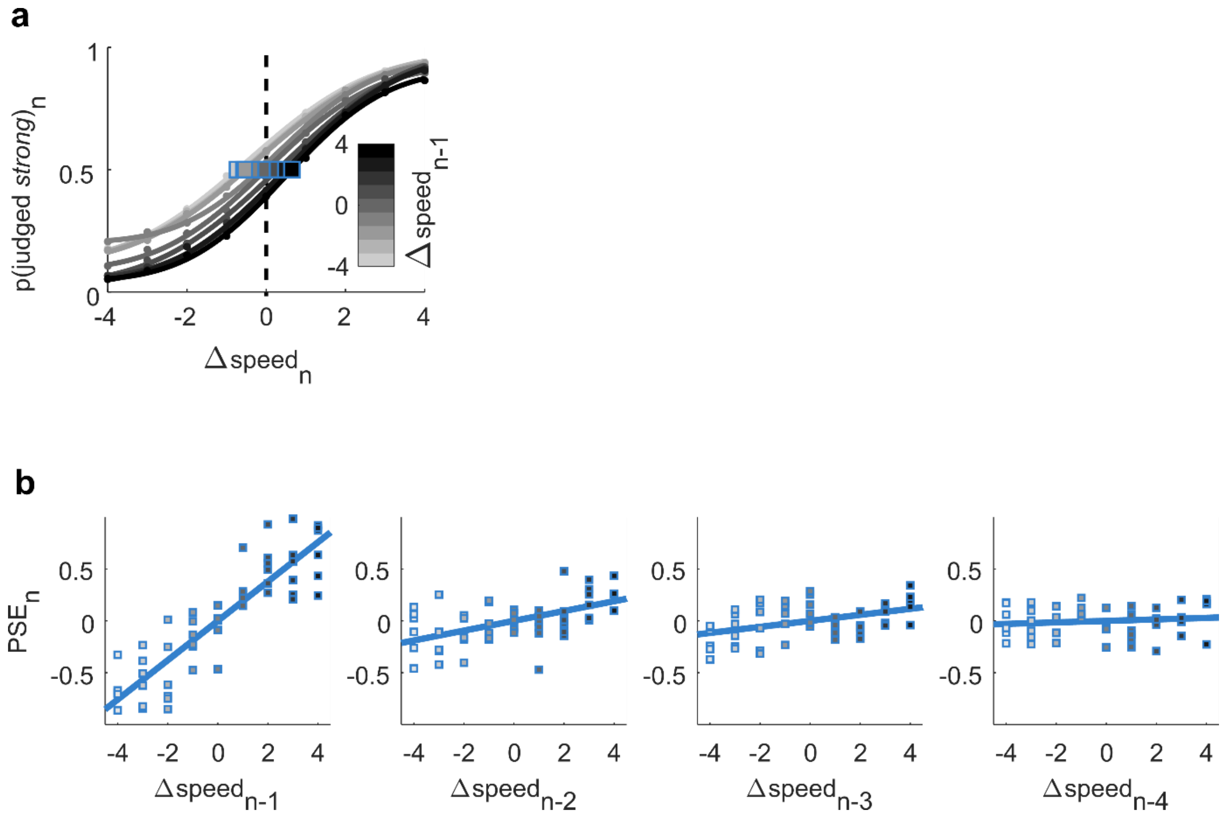


**Supplementary Figure S1. Gradual psychometric shift, excluding trials with incorrect *n-1* choice, suggests that *n-1* Δ*speed* drives the repulsive bias. a.** Probability of categorizing stimulus *n* as “strong” as a function of trial *n* Δ*speed*, with curves grouped by Δ*speed* of trial *n-1*, only using correct choices in trial *n-1*. Darker curves correspond to higher *n-1* Δ*speed*. Blue squares denote PSE. **b.** Bias of the trial *n* psychometric curve, depending on Δ*speed* in trial *n-1* (far left plot) to trial *n-4* (far right plot), after removing incorrect trials in the corresponding trial *n-1* to trial *n-1*. Shading of the squares denotes Δ*speed* in trial *n-1*. Squares correspond to individual rats.

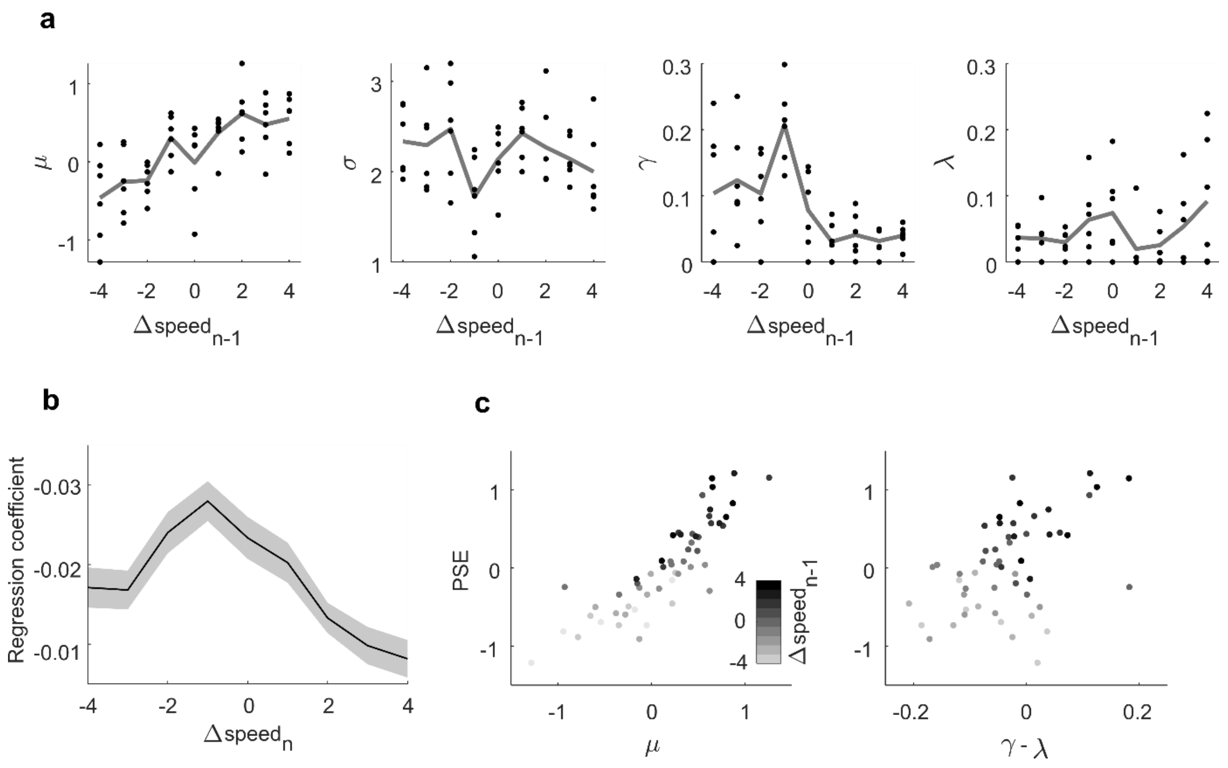


**Supplementary Figure S2. The effect of stimulus *n-1* on psychometric parameters and classification of stimulus *n* indicates horizontal psychometric shift. a.** Effect of Δ*speed* in trial *n-1* on psychometric curve parameters (curves, see Fig. 3a). Lower lapse rate was marginally modulated as a function of trial *n-1* Δ*speed* ($p=0.036$), while slope and upper lapse rate were not. The parameter most strongly influenced by previous stimulus was the curve midpoint ($p<0.001$), confirming horizontal shift as the psychometric curve property most influenced by the previous trial. **b.** Regression slopes for p(judged "strong") X Δ*speed* in trial *n-1*, for each trial *n* stimulus value. The resulting "bell" shape, with greater regression slopes for values of Δ*speed* near 0 indicates a greater effect of the previous stimulus on the evaluation of central Δ*speed* than on extreme Δ*speed*. Transparent shading represents standard deviation of the bootstrapped regression coefficients. **c.** Change in PSE as a function of considering only previous correct trials (lapses may change depending on previous error) curve midpoint μ (left panel, R2 = 0.7422) and PSE versus combined lapse rate γ–λ (right panel, R2 = 0.2242). Gray scale according to Δ*speed* in trial *n-1*. Only previous correct trials were considered (lapses may change depending on previous error).

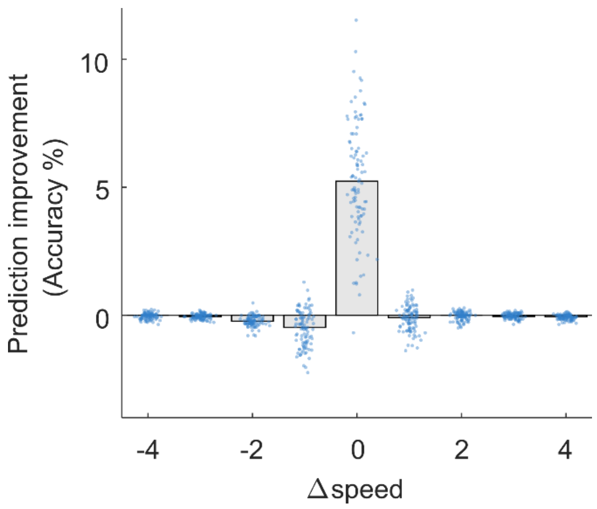


**Supplementary Figure S3. Discrete history model improves prediction about judgment of ambiguous stimuli only.** Improvement in performance of the discretized history-dependent model with respect to the no-history model (ideal observer) in predicting responses to each of the nine stimuli. Blue dots are cross-validation rounds over rats and sessions, grey bars illustrate the median.

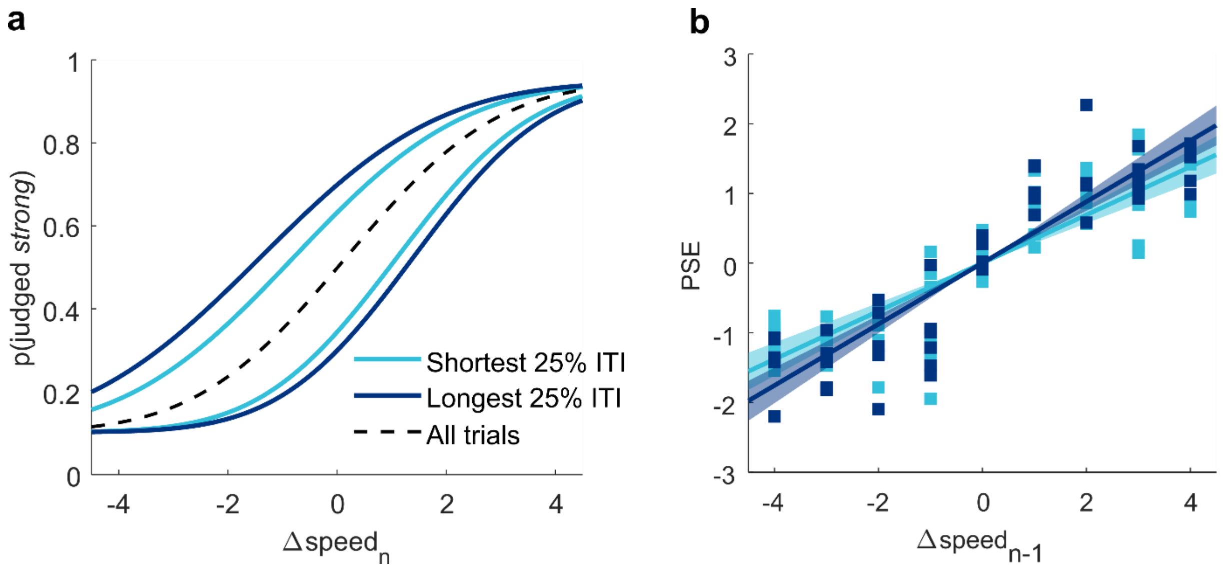


**Supplementary Figure S4. Stimulus-dependent bias builds up between trials also when removing incorrect trials and considering only early session trials. a.** Psychometric curves on trial *n* plotted according Δ*speed* of stimulus *n-1* and ITI, only using correct choices in trial *n-1* and early session trials (n<150; loss of motivation towards the end of the session might affect ITI duration). Dashed line is the average psychometric curve for all trials. **b.** Slope of the regression line fit between bias (PSE) and previous trial Δ*speed* for all rats, separated for shortest 2 ITI quartiles (light blue) and longest 2 ITI quartiles (dark blue), as in Figure 5c. Shading represents 95% confidence intervals. All trials with incorrect choices in trial *n-1* were removed and only early session trials were considered (see above). (interaction term between previous Δ*speed* and ITI durations, p=0.03).

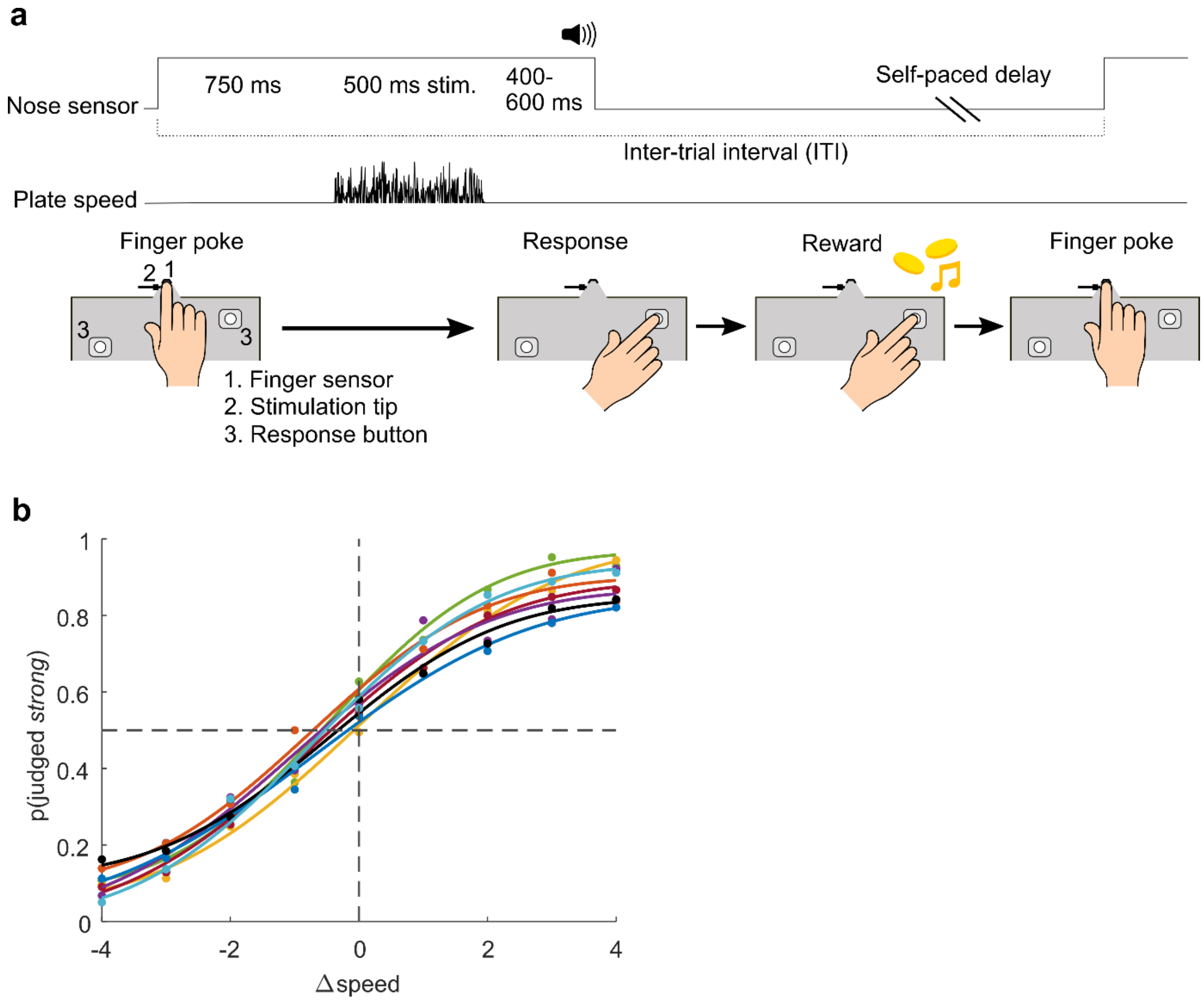


**Supplementary Figure S5. Vibrotactile categorization task transferred to human tactile perception. a.** Trial configuration. By placing her/his right index finger in the finger poke, the subject triggered a stimulation delivered to the fingertip by means of a vibrating rounded probe. The subject withdrew upon the go cue, and signaled her/his choice side by pressing a button either on the left or the right side. Buttons were asymmetrically placed in order to minimize motor biases (rule was switched between testing sessions). Subjects received feedback (correct/incorrect) on each trial through a computer monitor and headphones. **b.** Psychometric functions fitted to the averaged data of 8 human subjects (approximately 1,500 trials per subject).
